# A small epitope shared by p53 and an unrelated protein upregulated after adenovirus infection

**DOI:** 10.1101/2023.04.28.538733

**Authors:** Jessica J. Miciak, Fred Bunz

**Affiliations:** Department of Radiation Oncology and Molecular Radiation Sciences, Sidney Kimmel Comprehensive Cancer Center, and the Cellular and Molecular Medicine Graduate Program, The Johns Hopkins University School of Medicine, Baltimore, Maryland, USA

**Author notes:** Corresponding author: Fred Bunz, 1550 Orleans Street, Room 453, Baltimore, MD 21287.

## Abstract

Cancers commonly harbor point mutations in *TP53* that cause overexpression of functionally inactive p53 proteins. These mutant forms of p53 are immunogenic, and therefore present tantalizing targets for new forms of immunotherapy. Understanding how the immune system recognizes p53 is an important prerequisite for the development of targeted therapeutic strategies designed to exploit this common neoantigen. Monoclonal antibodies have been extensively used to probe the structural conformation of the varied isoforms of p53 and their respective mutants, and are still indispensable tools for studying the complex biological functions of these proteins. In this report, we describe the mapping of a novel epitope on p53 that appears to be shared by heat shock proteins (HSPs), which are typically upregulated in response to a variety of viral infections.

## Introduction

The signaling pathways mediated by the tumor suppressor p53 are altered in a majority of human cancers (1). The most frequent alterations are inactivating point mutations in the *TP53* exons that encode the central DNA binding domain. These mutations typically prevent p53 from productively binding to its cognate response elements throughout the genome and thus cause a loss of p53-dependent gene expression. In addition, the mutations that inactivate the transcriptional function of p53 also disrupt its interactions with E3 ubiquitin ligases, primarily MDM2, that normally keep the steady state levels of p53 low. For this reason, cancer cells that harbor inactivating p53 mutations will commonly express elevated levels of p53 protein compared with normal cells.

Despite its intracellular location, p53 protein is known to be immunogenic. Indeed, it was originally discovered as what we would now call a cancer-associated neoantigen (2), and detected by immunologic methods by virtue of its physical association with viral proteins (3). The first structural analyses of p53 were conducted with monoclonal antibody probes, which were able to distinguish mutant and wild type conformations of the protein (4). Subsequently, antibodies against p53 were widely used for the immunohistochemical analysis of tumor specimens to indirectly ascertain their *TP53* mutation status, prior to the era of high-throughput DNA sequencing.

The *TP53* locus expresses a number of proteins in addition to the full-length protein that has been most thoroughly studied (5). Two proteins lack the first 40 or 133 amino acid residues at the N-terminus, and are respectively known as Δ40p53 and Δ133p53. These p53 isoforms lack the major binding site for MDM2 and are therefore alternatively regulated. As a practical matter, Δ40p53 and Δ133p53 cannot be detected with the most commonly used anti-p53 monoclonal antibodies because these standard agents recognize an N-terminal epitope that is only present in the full-length isoform.

Adenoviruses express a protein called E1B-55K that binds p53 and targets it for degradation (6). This is one mechanism of several that this particular virus uses to suppress p53 activity. Large-scale studies of the human virome have revealed the remarkable diversity of viral proteins that directly bind p53, underscoring its importance as a host restriction factor against viral infection (7).

We were initially interested in exploring whether the smaller p53 isoforms might be similarly regulated by E1B-55K or other viral proteins. While probing cell lysates with a commercially-available but relatively obscure mouse monoclonal anti-p53 antibody designated 5H7B9, we found that it cross-reacted with 70 kDa host proteins that were robustly induced after infection. Peptide mapping revealed an epitope that is likely shared by p53 and HSP70 proteins, a family of molecular chaperones that are strongly upregulated in response to viral infection and a variety of other stressors.

## Results and Discussion

Full-length p53 is upregulated in response to DNA damage, but degraded following infection with wild type human adenovirus serotype 5 (Ad5). To examine the regulation of other p53 isoforms in response to these agents, we treated hTERT-RPE1 cells with doxorubicin, a chemotherapeutic agent that causes DNA damage, or infected them with wild type Ad5 at a high multiplicity of infection (MOI).

The widely used anti-p53 monoclonal antibodies DO-1 and DO-7 both recognize a compact epitope located between residues S20 and L25 (8). To permit an analysis of the p53 isoforms that lack this domain we used a commercially available mouse monoclonal antibody that was purported by the manufacturer (GenScript, see Materials and Methods) to recognize all known p53 isoforms.

Protein bands corresponding to full-length p53 (FLp53), Δ40p53 and Δ133p53 were clearly detectable (Fig 1A). The level of FLp53 was increased after doxorubicin treatment, as expected, while the abundance of the remaining p53 isoforms appeared unchanged. Notably, a major band migrating near the 70 kDa marker was observed in the Ad5-infected sample. Considering the prominence of this band and is exclusive appearance in cells infected with Ad5, we decided to investigate further.

**Figure 1.**
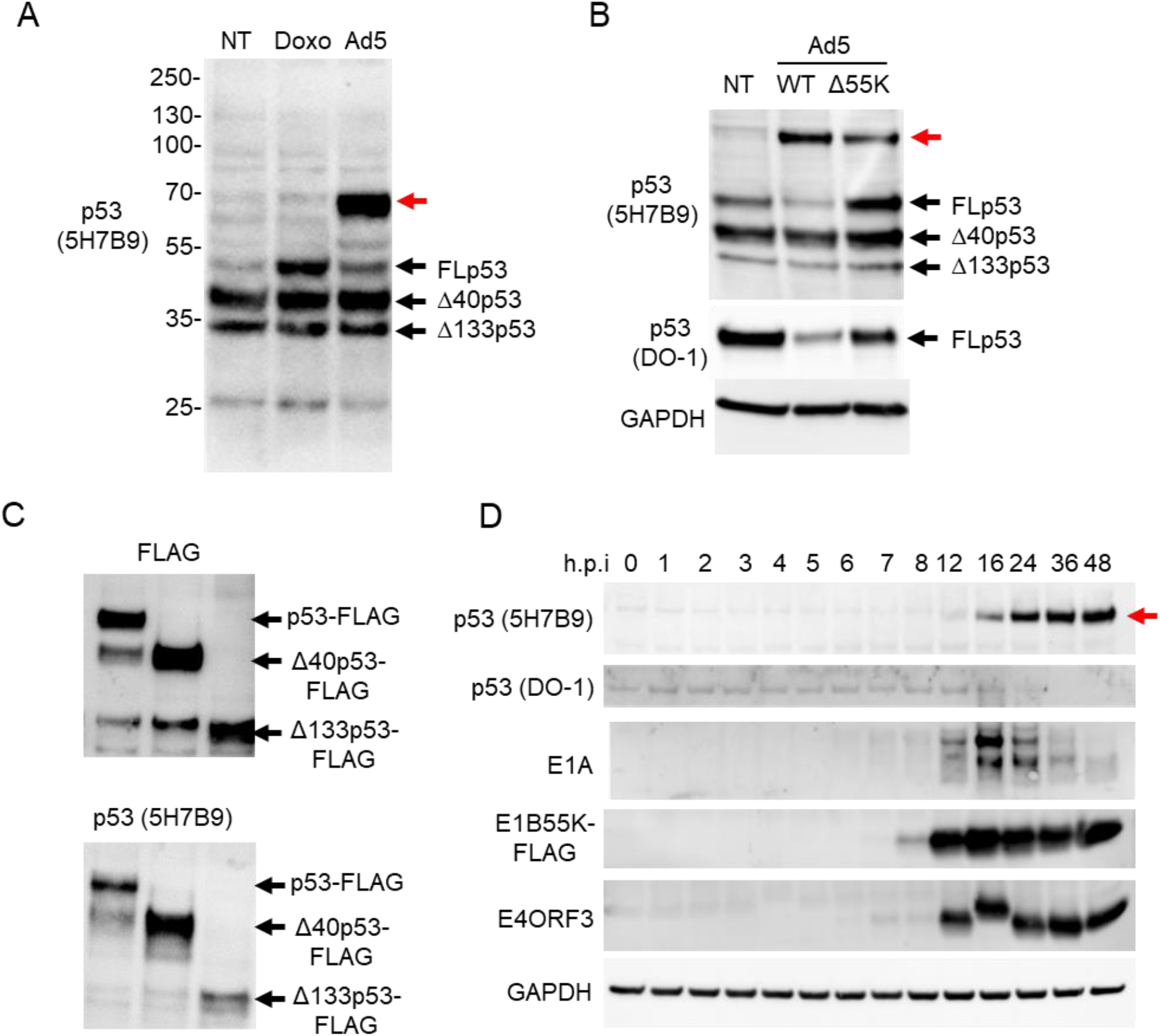
A 70 kDa protein induced after adenovirus infection cross reacts with a monoclonal antibody raised against p53. ***A***, hTERT-RPE1 cells were untreated (NT) or treated with 1μM doxorubicin (Doxo) or Ad5 (MOI=100) for 16 h. Protein extracts were probed with the mouse monoclonal antibody p53-5H7B9. The p53 isoforms are labeled, the 70 kDa band of interest is indicated by the red arrow. ***B***, Lysates prepared 16 h after infection with wild type Ad5 (WT) or a mutant that does not express the E1B-55K protein (Δ55K) were probed with the anti-p53 antibodies 5H7B9 and DO-1. GAPDH was detected as a loading control. ***C***, H1299 cells were transiently transfected with expression constructs encoding p53 isoforms with N-terminal FLAG tags. Lysates were collected 24 h following transfection and probed with an antibody against FLAG or p53-5H7B9. ***D***, hTERT-RPE1 cells were harvested at various time points following infection with a E1B-55K-FLAG adenovirus. Cell lysates were probed for the indicated proteins. Hours post-infection, h.p.i.

The level of FLp53, as assessed with the 5H7B9 antibody, was reduced following infection with wild type Ad5 and somewhat increased following infection with a mutant Ad5 that was defective for the expression of E1B-55K (Fig 1B). These results were consistent with the known function of E1B-55K, and further validated by detection of p53 with the DO-1 antibody, which only recognizes the full-length protein. In contrast, the unknown ∼70 kDa protein was induced by both viruses.

To further confirm that the 5H7B9 antibody was in fact recognizing the various isoforms of p53, we overexpressed FLAG-tagged versions of FLp53, Δ40p53 and Δ133p53 in H1299 cells, which express no endogenous p53 proteins (Fig 1C). Similar bands were detected by anti-FLAG antibody and 5H7B9, confirming the specificity of 5H7B9 for p53.

We next infected cells with a recombinant Ad5 virus in which a FLAG epitope tag had been inserted at the end of the E1B-55K coding sequence (Ad5-E1B-55K-FLAG (9)), allowing us to track this critical viral protein. The cross-reactive protein(s) appeared within 16 h of infection with Ad5-E1B-55K-FLAG (Fig 1D), shortly following the expression of the early viral proteins E1A and E4ORF3. The E1B-55K-FLAG protein could be detected by 8 h after infection, and was maximally expressed 4 h later.

These experiments demonstrated that an as-yet unidentified ∼70 kDa protein could be robustly and reproducibly induced following adenovirus infection, and subsequently detected with an antibody that had been raised against p53 and which recognized the three major p53 isoforms. To gain further insight into the relationship between this cross-reactive protein and the adenovirus life cycle, we incubated the p53-5H7B9 antibody with hTERT-RPE1 cells that had been infected with the Ad5-E1B-55K-FLAG virus, and detected its distribution by immunofluorescence (Fig 2).

**Figure 2.**
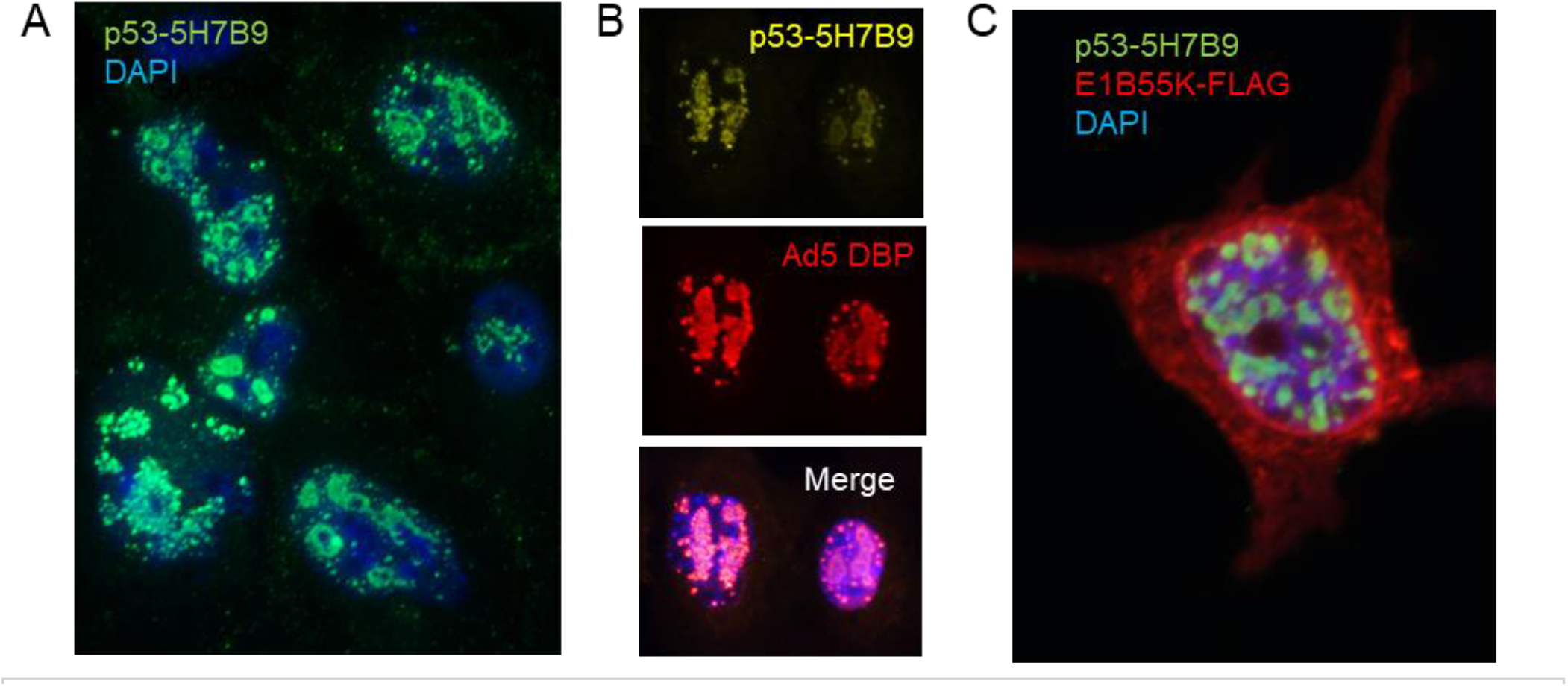
The 70-kDa cross-reactive protein localizes to adenovirus replication centers. ***A***, hTERT-RPE1 cells were infected with Ad5-E1B-55K-FLAG (MOI 10), for 24 h, fixed, permeabilized and stained with p53-5H7B9 antibody (Green). Nuclei were counterstained with 4’,6-diamidino-2-phenylindole (DAPI). ***B***, cells treated as in *A* were co-stained with p53-5H7B9 (green) and an antibody to the adenoviral DNA binding protein (Ad5-DBP, red). ***C***, cells treated as in ***A*** were co-stained with p53-5H7B9 (green) and anti-FLAG (Red). The nucleus was counterstained with DAPI.

Staining with p53-5H7B9 revealed an intricate, elliptical pattern of nuclear localization (Fig 2A) that is highly characteristic of adenoviral replication compartments (10). This interpretation was supported by co-localization of the 5H7B9 signal and the detection of Ad5 DNA binding protein, a core component of viral DNA replication complexes. The E1B-55K-FLAG protein appeared to be primarily localized to the nuclear periphery, consistent with the role of E1B-55K in RNA export (11). We can infer that the detected protein was not p53, as p53 is rapidly degraded by the virus. The association between the p53-5H7B9-reactive protein and nuclear Ad5 replication centers suggested that the cross-reactive protein was not merely upregulated by the cellular stress caused by viral infection, but was instead directly involved in the adenovirus life cycle.

To identify this cross-reactive protein, we first mapped the precise 5H7B9 interaction site on p53. A peptide scanning array was designed to cover the Δ133p53 variant, which our previous experiments implicated as the minimal region of interest (Fig 1C). When probed with the 5H7B9 antibody, three spots gave a strong signal (Fig 3A). These spots corresponded to three overlapping peptides that commonly contained the p53 amino acid residues 264-274 (Fig 3B).

**Figure 3.**
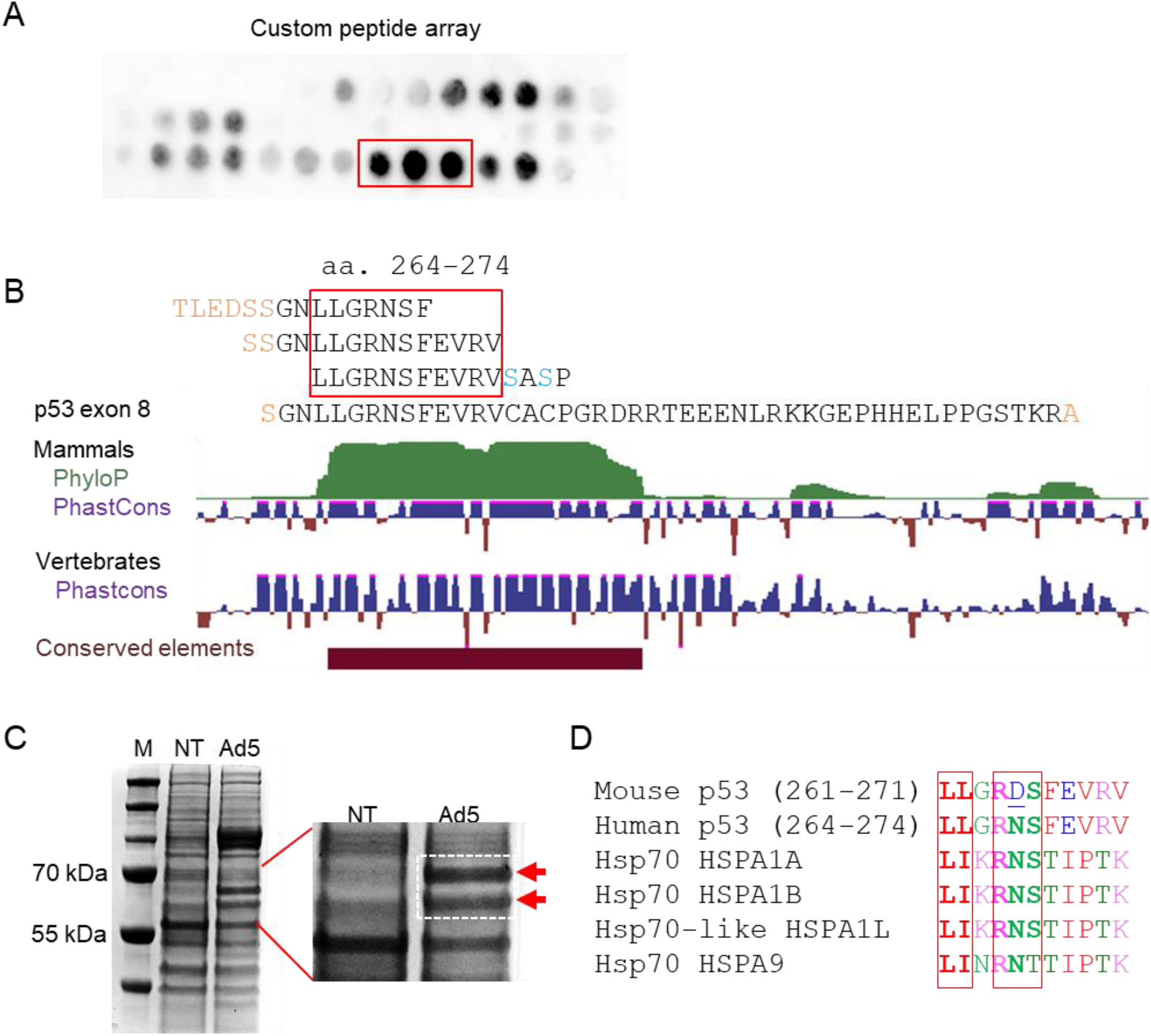
Mapping the p53-5H7B9 epitope by peptide scan. ***A***, A custom-designed PepSpot peptide array was probed with the p53-5H7B9 antibody, as described in Materials and Methods. The three spots with the highest signal above background (red box) corresponded to the three peptides shown in panel ***B***. The cysteine residues in the p53 sequence (shown in blue) are represented by serine in the peptide array. An illustration showing the evolutionary conservation in this region was rendered by the UCSC Genome Browser. ***C***, RIPA extracts from non-treated (NT) and Ad5 infected cells were resolved by 1D gel electrophoresis and stained with Coomassie Blue. The relevant region of the gel, defined by molecular weight markers (M) was cut out for proteomic analysis. Major bands specific to Ad5 infection are indicated by red arrows. ***D***, The proteins shown were aligned by Clustal Omega. The putative binding motif is boxed in red.

We next fractionated RIPA extracts from uninfected and Ad5-infected cells by 1D-PAGE (Fig 3C). Adenovirus inhibits host protein synthesis, and we accordingly observed an overall decrease in the proteins extracted from Ad5 infected cells. A gel slice encompassing the ∼65-75 kDa size range, previously defined by our western blots (Fig 1A), was excised. Proteins were subject to in-gel digestion with trypsin, and the resulting peptides were resolved by mass spectrometry.

A number of nuclear proteins were represented in the extract from Ad5-infected cells (see Table 1). Several of these hits, including RPA1, RECQ1, XRCC6, and DDX3X, mediate various aspects of nucleic acid metabolism. The latter three proteins are normally present at low abundance in unstressed cells. Each of these proteins could conceivably be involved in adenovirus DNA replication or gene expression, processes that are highly dependent on the recruitment of host proteins. Relatively abundant peptides extracted from this size fraction mapped to heat shock proteins. Ubiquitous molecular chaperones that function in folding and assembly of client proteins, the heat shock proteins play important roles during adenovirus infection and are strongly induced at the transcriptional and posttranscriptional levels (12, 13). Interestingly, peptides corresponding to adenovirus-encoded proteins were not detected with any significant level of confidence.

**Table 1.**
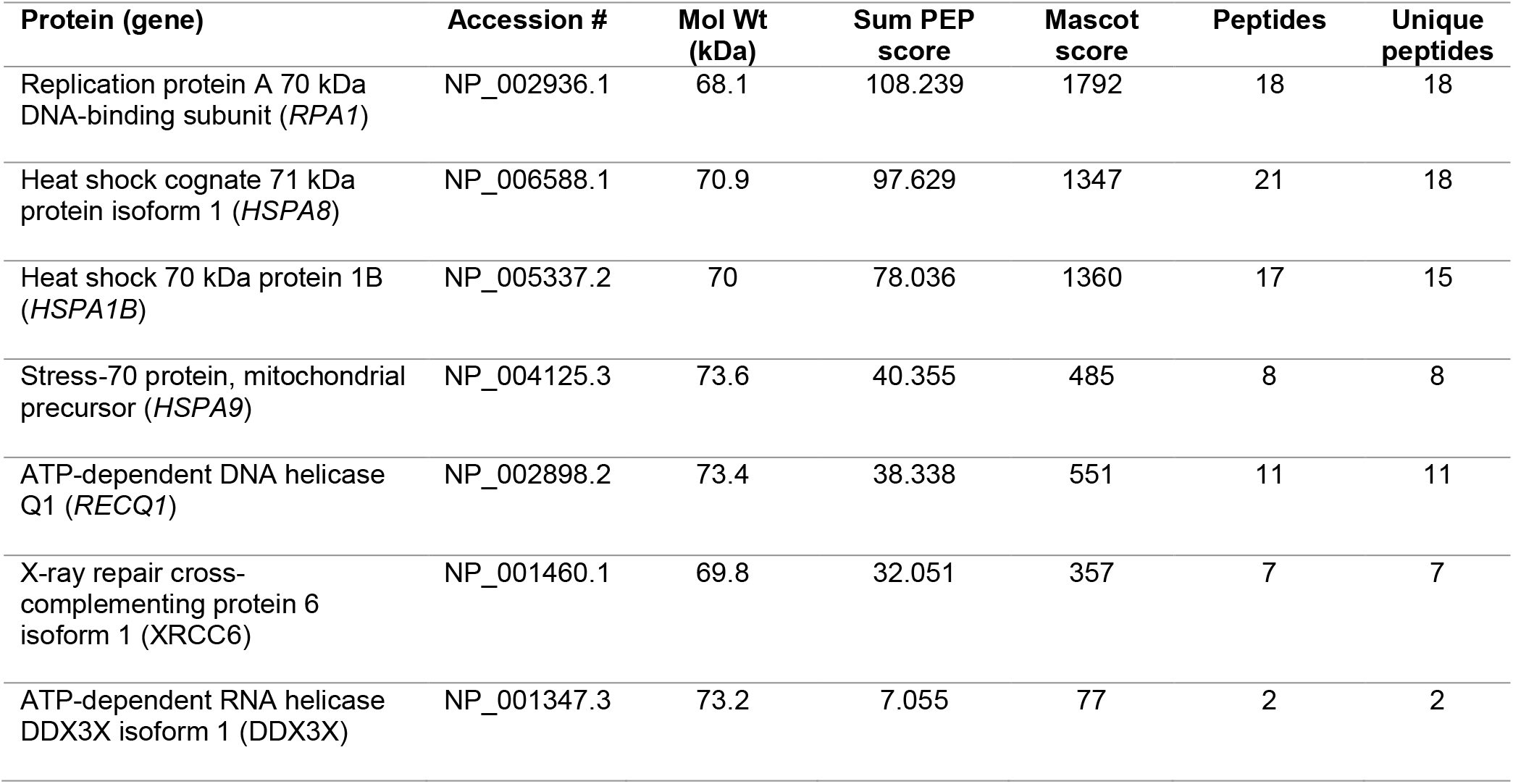
Selected proteins identified in Ad5-infected extracts. High abundance peptides mapping to keratins and proteins not of human origin were excluded from further analysis.

We systematically aligned the deduced sequence of the putative p53 epitope against the proteins identified in the infected cell lysate with Clustal Omega software. This analysis revealed a short region of homology shared by p53 and several heat shock proteins. Within the experimentally defined 11-amino acid region, the sequence motif LL/IXRNS was present in human p53, and also in the heat shock proteins encoded by *HSPA1A, HSPA1B* and *HSPA1L*. While this region is evolutionarily conserved (Fig 3B), murine p53 diverges at a single position within this motif (LL/IXR**<u>D</u>**S, Fig 3D). The p53-5H7B9 antibody failed to recognize any of the p53 proteins expressed in mouse NIH3T3 cells after irradiation (not shown), providing further support for this epitope as the precise binding site of the p53-5H7B9 antibody.

Together, these data suggest that the LL/IXRNS motif shared by p53 and several proteins in the Hsp70 family comprises a critical part of the p53-5H7B9 epitope. The well-described upregulation of these proteins in response to adenovirus infection would provide a plausible explanation for the induction of the cross-reactive band observed on western blots and the remarkable localization of this protein at viral replication centers. While our data generally support this conclusion, it is difficult to definitively rule out other proteins that might account for these observations.

Monoclonal antibodies raised in mice were critical reagents for the early characterization of p53, and remain important tools for understanding p53 biology. A simple search of the Pubmed database with the term ‘p53’ produces a list of more than 115,000 papers. It is safe to assume that p53-specific antibodies were employed in a large proportion of the primary studies of p53, and will continue to be mainstays in the field of p53 research.

The monoclonal antibodies DO-1 and DO-7, which are among the most useful and widely distributed reagents for detection of p53, recognize a common epitope near the N-terminus, and therefore cannot detect the isoforms that lack this region (Fig 4A). The N-terminal region of p53 is relatively unstructured (14); biophysical studies of p53 and its interactions with DNA have largely focused on a core domain that is more structurally stable (Fig 4B). There remains a relative paucity of commercially available antibodies that can recognize this central part of the protein, which is critical to p53 function.

**Figure 4.**
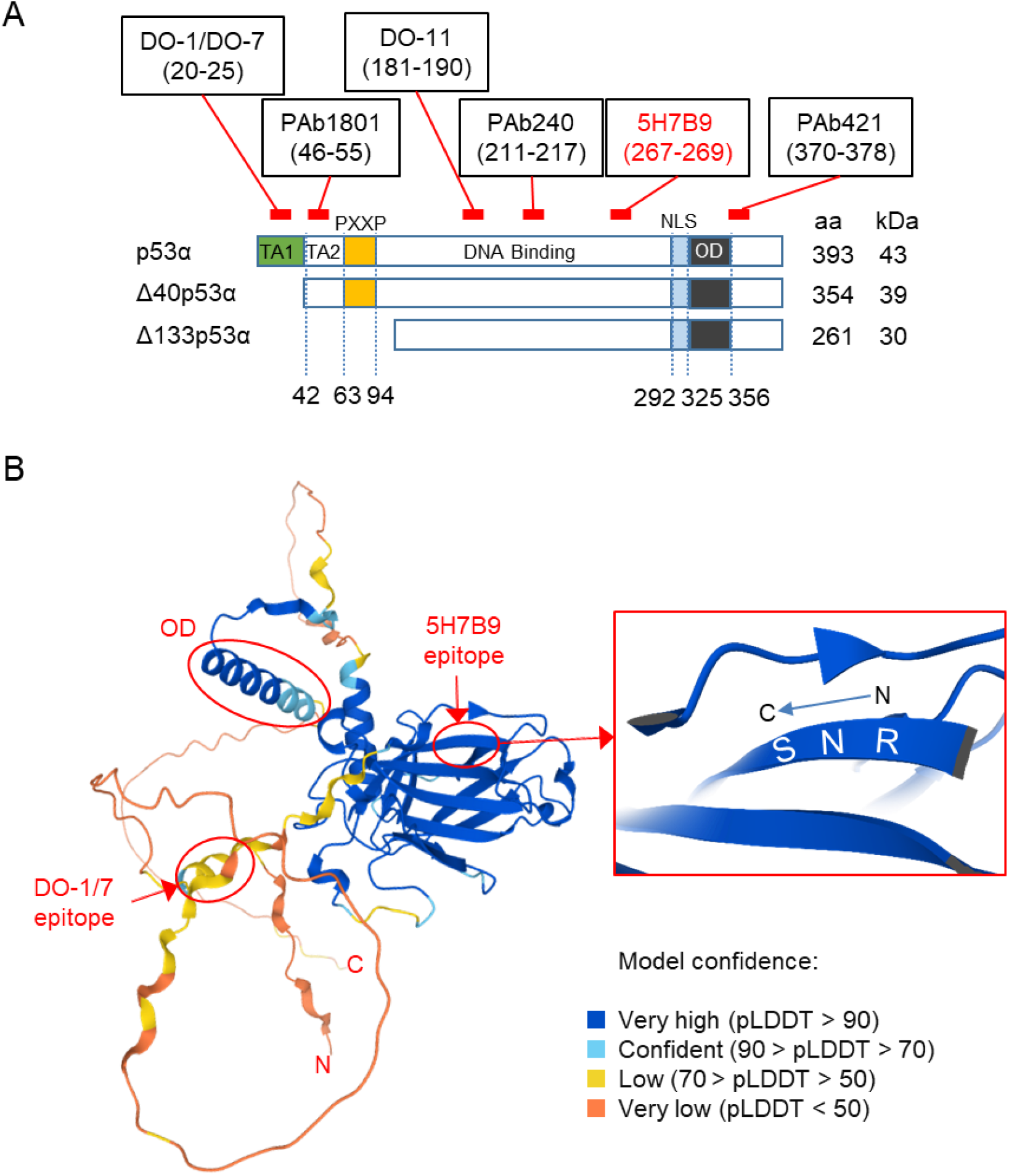
Useful antibodies against human p53. ***A***, A map shows the epitopes of widely distributed p53 antibodies with respect to the three major p53 isoforms and their functional domains. The transactivation domains (TA1,2), the proline-rich domain (PXXP), the DNA binding domain, the nuclear localization signal (NLS), and the c-terminal oligomerization domains are indicated. ***B***, An AlphaFold ribbon structure prediction based on 245 p53 structures in PDB linked to UniProt accession ID P04637. The defined epitopes for the DO-1/DO-7 and 5H7B9 antibodies are indicated. Oligomerization domain, OD.

This study has identified an epitope that can be used for the immunologic detection of full-length p53 as well as several relatively understudied isoforms that lack the N-terminus. The deduced minimal epitope, including amino acid positions 269-271 is infrequently altered by mutation, and would presumably be conserved in the so-called hotspot mutants, which affect R175, Y220, G245, R248, R273 and R282. We have not tested the ability of p53-5H7B9 to detect mutant proteins. The p53-5H7B9 antibody would presumably be unable to detect the recently described isoform p53Ψ, which is generated by alternative splicing leading to a frame-shift upstream of the mapped region (15).

Our study also shows that the use of the 5H7B9 antibody for the detection of p53 comes with the caveat that it may also recognize other important proteins in the cell stress responses. Indeed, many useful antibodies recognize cross-reactive species, but the protein sources of these spurious signals usually remain undefined. The overall significance, if any, of the shared p53-HSP70 epitope is unclear. It would seem improbable but not inconceivable that a single antibody could by chance recognize the same small epitope on two protein families that are inducible upon viral infection. As p53 increasingly becomes in focus as a target for new modes of immunotherapy, this study might serve as a useful reminder that relatively small structural homologies can lead to off-target effects.

The production of the p53-5H7B9 antibody by the commercial vendor had been suspended at the time of this writing, and the catalog presently lists this item as discontinued. It remains unclear if the hybridoma line is still available. If not, our study suggests that the peptide sequence identified as the p53-H7B9 epitope might in the future be used to generate useful new reagents for the detection of the multifunctional proteins expressed from the *TP53* gene.

## Materials and Methods

### Cell culture

The telomerase-immortalized human cell line hTERT-RPE1 was a gift from Prasad Jallepalli. Cells were maintained at 37°C in 5% CO_2_ in DMEM/F12 medium supplemented with 6% fetal bovine serum (FBS) and penicillin/streptomycin. These cells were authenticated by Short Tandem Repeat (STR) profiling and tested for the presence of mycoplasma at the Johns Hopkins Genomic Resources Core Facility.

### Antibodies

The p53-5H7B9 mouse monoclonal antibody (subclass IgG2a) was purchased from GenScript (catalog number A01767-40) as lyophilized protein in PBS, and resuspended in deionized water to a concentration of 0.5 mg/ml. According to the manufacturer, p53-5H7B9 was produced by a hybridoma resulting from fusion of SP3/O-Ag14 myeloma and B lymphocytes from a mouse immunized with full length recombinant p53 protein. Other antibodies used in the study included anti-p53 mouse monoclonal DO-1 and anti-E1A mouse monoclonal M58, both from Santa Cruz Biotechnology, anti-FLAG (DYKDDDDK) rabbit monoclonal D6W5B from Cell Signaling Technology, and anti-GAPDH rabbit polyclonal from Millipore Sigma. The anti-E4ORF3 antibody was a gift from Gary Ketner. The anti-Ad5-DBP antibody was a gift from David Ornelles.

### Western blots

Proteins were extracted in RIPA buffer (Cell Signaling Technologies), resolved on Bolt Bis-Tris mini-gels (ThermoFisher Scientific) and transferred to PVDF membranes (MilliporeSigma). The reconstituted p53-5H7B9 stock was diluted in TBST/5% bovine serum albumin to a final concentration of 0.5 µg/ml. The remaining antibodies were used at 1:1000 dilution. Primary antibodies were incubated with blocked membranes for 18-24 h at 4°C. After three washes in PBS, blots were developed with horseradish peroxidase-conjugated secondary antibodies followed by enhanced chemiluminescence.

### PepSpot screening array

A custom array composed of 64 individual 15-mer peptides with 11 amino acid overlaps was designed to cover the protein sequence between p53 amino acids 133-306. The peptides were synthesized and spotted on a PVDF filter by JPT Peptide Technologies. Each peptide was acetyl-blocked at the N-terminus. Cysteine residues were replaced by serine. The resulting array was probed with the p53-5H7B9 antibody using the conditions for western blotting, as described above.

### Immunofluorescence

Cells grown on chamber slides (Labtek) were fixed and permeabilized with freshly prepared 3% paraformaldehyde/0.1% TritonX-100 for 30 min at room temperature. Primary antibodies were used at 1:100 dilution.

### In-gel digestion and characterization of proteins induced by Ad5

Approximately 500 µg of protein in RIPA buffer were loaded on a 1D Bolt 4-12% Bis-Tris gel and resolved by electrophoresis. The gel was stained with Coomassie blue and imaged. Following the excision of the region of interest, the gel slice was washed with 50% methanol. The gel matrix was then dehydrated in 100% acetonitrile and reswollen with trypsin digestion solution on ice, then incubated at 37°C overnight. Tryptic peptides were extracted by dehydrating the gel in 80% acetonitrile. Extracted peptides were dried by vacuum centrifugation, resuspended in 0.1% formic acid and analyzed by liquid chromatography interfaced with tandem mass spectrometry. Fragmentation spectra were searched against the protein database with an auto-concatenated decoy database to identify peptide sequence and protein at a confidence of 5% False Discovery Rate. This proteomic analysis was performed at the Sidney Kimmel Comprehensive Cancer Center Mass Spectrometry and Proteomics Core

## Acknowledgements

The authors are grateful to our colleagues for generously providing critical reagents. These studies were supported by the NIGMS (R01GM135485), the NCI (P30CA006973) and the Emerson Collective.

